# HIV reservoirs are dominated by genetically younger and clonally enriched proviruses

**DOI:** 10.1101/2023.04.12.536611

**Authors:** Natalie N. Kinloch, Aniqa Shahid, Winnie Dong, Don Kirkby, Bradley R. Jones, Charlotte J. Beelen, Daniel MacMillan, Guinevere Q. Lee, Talia M. Mota, Hanwei Sudderuddin, Evan Barad, Marianne Harris, Chanson J. Brumme, R. Brad Jones, Mark A. Brockman, Jeffrey B. Joy, Zabrina L. Brumme

## Abstract

In order to cure HIV, we need to better understand the within-host evolutionary origins of the small reservoir of genome-intact proviruses that persists within infected cells during antiretroviral therapy (ART). Most prior studies on reservoir evolutionary dynamics however did not discriminate genome-intact proviruses from the vast background of defective ones. We reconstructed within-host pre-ART HIV evolutionary histories in six individuals and leveraged this information to infer the ages of intact and defective proviruses sampled after an average >9 years on ART, along with the ages of rebound and low-level/isolated viremia occurring during this time. We observed that the longest-lived proviruses persisting on ART were exclusively defective, usually due to large deletions. In contrast, intact proviruses and rebound HIV exclusively dated to the years immediately preceding ART. These observations are consistent with genome-intact proviruses having shorter lifespans, likely due to the cumulative risk of elimination following viral reactivation and protein production. Consistent with this, intact proviruses (and those with packaging signal defects) were three times more likely to be genetically identical compared to other proviral types, highlighting clonal expansion as particularly important in ensuring their survival. By contrast, low-level/isolated viremia sequences were genetically heterogeneous and sometimes ancestral, where viremia may have originated from defective proviruses. Results reveal that the HIV reservoir is dominated by clonally-enriched and genetically younger sequences that date to the untreated infection period when viral populations had been under within-host selection pressures for the longest duration. Knowledge of these qualities may help focus strategies for reservoir elimination.

**Importance:** Characterizing the HIV reservoir that endures despite antiretroviral therapy (ART) is critical to cure efforts. Our observation that the oldest proviruses persisting during ART were exclusively defective, while intact proviruses (and rebound HIV) all dated to the years immediately pre-ART, explains why prior studies that sampled sub-genomic proviruses on-ART (which are largely defective) routinely found sequences dating to early infection, whereas those that sampled viral outgrowth sequences found essentially none. Together with our findings that intact proviruses were also more likely to be clonal, and that on-ART low-level/isolated viremia originated from proviruses of varying ages (including possibly defective ones), our observations indicate that: 1) on-ART and rebound viremia can have distinct within-host origins, 2) intact proviruses have shorter lifespans than grossly-defective ones, and therefore depend on clonal expansion for persistence, and 3) the HIV reservoir, being overall genetically younger, will be substantially adapted to within-host pressures, complicating immune-based cure strategies.

## Introduction

Following infection, Human Immunodeficiency Virus 1 (HIV-1) integrates its genome into that of the host cell, usually a CD4+ T-lymphocyte^1, 2^. Most infected cells die, or are eliminated by the immune system^3^, usually within two days of infection^4^, but a minority persist even during long-term antiretroviral therapy (ART), and can fuel viral rebound if ART is interrupted^5–8^. If we are to cure HIV, it is critical to understand the within-host evolutionary origins and dynamics of the genome-intact and replication-competent proviruses that comprise this persistent HIV reservoir.

Seeding of HIV sequences into the reservoir begins immediately following infection^9–12^, and continues until ART initiation^13–16^. During untreated infection however, turnover is relatively rapid^14, 16–18^, where recent half-life estimates are on the order of half a year, compared to nearly four years for intact proviruses during the initial years of ART^19–27^. As such, if ART is not initiated until chronic infection, most early within-host HIV lineages will have already been eliminated by this time, as demonstrated by the observation that most proviruses sampled during ART “date” to the year or two prior to ART initiation^13–16, 28, 29^. Nevertheless, older proviruses, some dating as far back as transmission, are also routinely recovered during ART^13–16^. While it is now becoming clear that host cell features, such as genomic integration site^30–37^ and clonal expansion^21, 31, 33, 38–47^ influence how long a HIV provirus will persist within-host, the contribution of viral genetic features to proviral persistence remains incompletely understood.

Strong evidence nevertheless supports such a relationship. The small pool of genetically intact proviruses that comprises the HIV reservoir, but that represents only ∼5% of all proviruses persisting on ART^48–50^, decays more rapidly during ART than the much larger pool of defective proviruses that harbor large deletions, hypermutation, packaging signal region defects, point mutations and/or other defects^21, 22, 25–27, 38, 48, 50^. In individuals who initiated ART in chronic infection therefore, the very long-lived proviruses that are routinely recovered during ART (*i.e.* those dating back to early infection) would thus be predicted to be defective, whereas intact proviruses should generally be “younger” (*i.e.* date to the year or two before ART initiation). No studies to our knowledge have leveraged information from pre-ART within-host evolutionary histories to simultaneously elucidate the integration dates of both intact and defective proviruses sampled during ART^13–16, 28, 29^: all but one prior study collected sub-genomic sequences only, which meant that they could not distinguish intact from defective proviruses^48, 50^, while the remaining study exclusively sampled replication-competent HIV following *ex vivo* stimulation, and therefore could not investigate age differences between intact and defective sequences^29^.

Our knowledge of the within-host evolutionary origins of HIV sequences reactivated from the reservoir *in vivo* − namely, HIV RNA sequences rebounding in plasma following ART interruption, as well as low-level and/or isolated viremia occurring during otherwise suppressive ART − also remains limited. While rebound virus likely originates from intact HIV proviruses, there is evidence that on-ART viremia can originate from both genome-intact^51^ and defective proviruses^52^. Indeed, as we now appreciate that proviruses with genomic defects can in some cases produce HIV transcripts, proteins^49, 53^ and even virions^52^, and that cells harboring them can be recognized by the immune system^53, 54^, achieving a deeper understanding of their longevity and potential to contribute to on-ART viremia is also important. To address these knowledge gaps, we employed a phylogenetic approach^13^ to reconstruct within-host HIV evolutionary histories and investigate the age distribution of intact proviruses, different types of defective proviruses, along with *in vivo* and *ex vivo* reactivated HIV sequences, in six individuals living with HIV who had been receiving ART for a median of more than 9 years.

## Results

### Participant characteristics and reservoir sampling

We isolated 2,336 near-full-length proviruses (median 352; range 195-733 per participant) at a single time point from six participants living with HIV who had been receiving ART for a median 9.3 (range 7.2- 12.2) years (**Table 1; Figure 1**). In contrast to some previous studies^21, 23, 40^, we used a strict definition of genome-intact that required all HIV proteins (including accessory proteins) to be intact. Also, as we retained only amplicons that were sequenced end-to-end (see methods), we could definitively classify each provirus as intact or defective (*i.e.* there was no “inferred intact” category). For four participants, we performed additional on-ART sampling as follows: from BC-003 and BC-004, we isolated 2 and 7 full-genome HIV RNA sequences, respectively, from limiting-dilution Quantitative Viral Outgrowth Assays (QVOA) performed on the same sample as used for proviral amplification. From BC-001 we isolated 9 subgenomic (*nef*) HIV RNA sequences from plasma after ART interruption. From BC-001, BC-002 and BC-004 we isolated 104 HIV RNA sequences from low-level or isolated viremia events during otherwise suppressive ART. We defined these as isolated or intermittent viremia generally below 1,000 HIV RNA copies/mL according to WHO guidelines^55^, though participants 2 and 4 had isolated measurements exceeding this, but with no record of ART interruption nor appearance of new antiretroviral resistance mutations.

**Figure 1:**
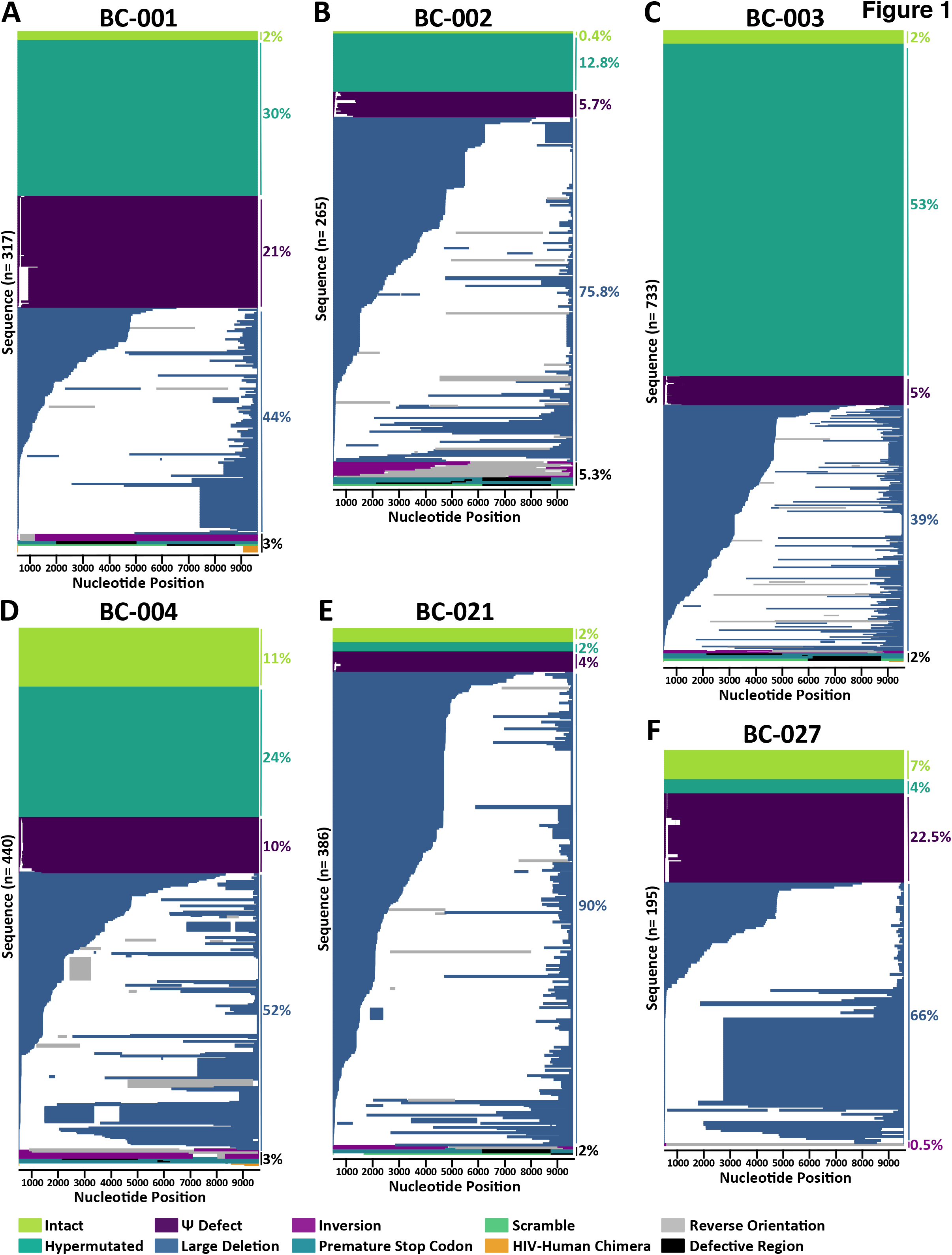
On-ART proviral landscapes. Proviral genomes isolated from BC-001 (n=317; *panel A*), BC-002 (n=265; *panel B*), BC-003 (n =733; *panel C*), BC-004 (n=440; *panel D*), BC-021 (n=386; *panel E*), and BC-027 (n=195; *panel F*), colored based on genomic integrity as indicated. The frequency of each proviral category is shown to the right. For defective sequences, white regions denote deletions and grey regions denote HIV regions that are in the reverse orientation. For the “premature stop codon” and “scramble” categories, black regions denote the region(s) containing the defect(s).

**Table 1:** Participant clinical history and HIV sequence sampling.

| Participant | Years from estimated infection to first ART<sup>a</sup> | Years of pre-ART sampling (n time points) | Number of pre-ART HIV RNA sequences collected (% unique) | Subsequent years of ART | Total number of proviral sequences collected on ART (% unique) | Number of intact proviruses collected on ART<sup>b</sup> (% unique) | Number of QVOA sequences collected on ART (% unique) | Number of rebound HIV RNA sequences collected (% unique) | Number of low-level viremia sequences collected (% unique) |
| --- | --- | --- | --- | --- | --- | --- | --- | --- | --- |
| BC-001 | 11.9 | 10 (15) | 102 (91%) | 9.5 | 317 (65%) | 5 (100%) | - | 9 (56%) | 1 (100%) |
| BC-002 | 14.6 | 9.5 (17) | 160 (82%) | 9.6 | 265 (75%) | 1 (100%) | - | - | 13 (77%) |
| BC-003 | 5.9 | 4.75 (8) | 122 (93%) | 8.7 | 733 (88%) | 15 (87%) | 2 (100%) | - | - |
| BC-004 | 2.2 | 0.75 (4) | 65 (91%) | 9.1 | 440 (69%) | 47 (53%) | 7 (100%) | - | 90 (9%) |
| BC-021 | 5 | 2.5 (8) | 221 (79%) | 12.2 | 386 (72%) | 9 (44%) | - | - | - |
| BC-027 | 26.5 | 14.25 (7) | 215 (82%) | 7.2 | 195 (54%) | 14 (43%) | - | - | - |

<sup>a</sup>Infection date was estimated using clinically-recorded information, participant-reported date or the mean root date of within-host phylogenies, whichever was earlier. 'First ART' refers to first treatment with triple ART
<sup>b</sup>This number is a subset of the total number of proviruses collected on ART

We also isolated 885 HIV RNA *nef* sequences (median 141; range 65-221 per participant) from a median 8 (range 4- 17) longitudinal archived plasma samples that spanned a median 7.1 (range 0.75- 14.25) years prior to ART. These sequences were used to reconstruct participants’ pre-ART HIV evolutionary histories, which were in turn used to infer the ages of HIV sequences persisting on ART. We used *nef* because it evolves rapidly within-host, but is nevertheless representative of within-host HIV diversity elsewhere in the genome^13^. Using *nef* also allowed us to maximize the number of sequences whose integration dates could be inferred, as it is the most likely region to be intact in proviruses persisting on ART^50^. All participants had HIV subtype B, and within-host sequences were monophyletic with no evidence of super- infection (**Supplemental Figures 1, 2**).

**Figure 2:**
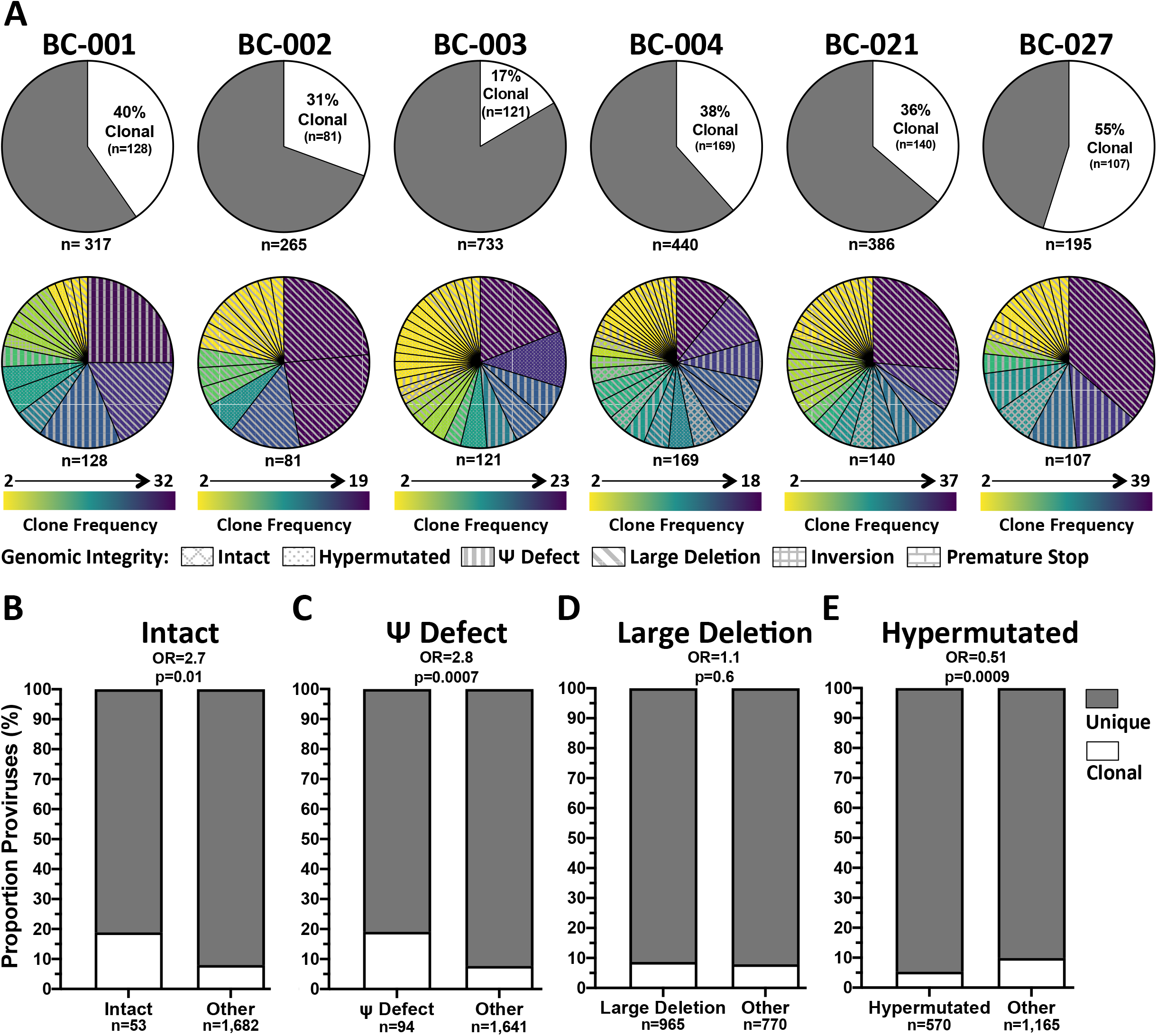
Proviral clonality and rela1onship with genomic integrity. *Panel A: (top)* White pie slices denote each participant’s proportion of clonal sequences (defined as 100% identical sequences observed at least twice). *(bo4om)* Breakdown of clonal sequences by clone size (color gradient) and genomic integrity (hatching). *Panels B-E:* Pairwise comparisons of the proportion of clonal sequences in each proviral category as indicated (clonal proportion shown in white), compared to all other provirus types. Data are combined across all participants. P- values are computed using Fisher’s exact test and are not corrected for mulNple comparisons.

### Proviral Landscape During Long-term ART

Of the 2,336 near full-length proviruses collected, only 4% were genome-intact: a median of 2% of sequences (range 0.4%- 11%; 1- 47 sequences) per participant (**Figure 1, Table 1, Supplemental Table 1**). BC-004 had the highest proportion of intact sequences, at 11% (n=47; 25 unique), while BC-002 had the lowest, at 0.4% (n=1) (**Figure 1B, 1D, Table 1, Supplemental Table 1**). For four participants (BC-001, BC-002, BC-003, BC-021), the recovered proportion of intact proviruses was lower than that predicted by the Intact Proviral DNA Assay (**Supplemental Table 1**), which is not surprising given the inefficiency of long-range PCR^56^ and the ability of sequencing to capture defects outside of the IPDA target regions^50^. Proviruses with large deletions dominated in all participants except BC-003, and made up a median 59%, range 39- 90% of proviruses/participant (**Figure 1A-F, Supplemental Table 1**). These varied greatly in length − the shortest, recovered from BC-004, was just 167 base pairs − and many had additional defects such as gene inversions, scrambles and hypermutation. We also observed evidence of template switching between repeated genomic elements during reverse transcription as a reproducible mechanism for large deletion formation, both within and between individuals. Thirteen distinct sequences from BC-003 for example had a deletion spanning HIV genomic nucleotides 4,781 - 9,064 (numbering according to the HXB2 reference strain), where the sequence flanking the deletion was ‘TTTTAAAAGAAAAGGGGGGA’. These exact breakpoints, which have been described by others^38, 48^, were also observed in one of BC-001’s proviruses, one of BC-004’s, and eleven distinct proviruses from BC-021.

By contrast, hypermutation dominated BC-003’s proviral landscape, at 53%. Prior CD4+ T-cell phenotyping of this participant had revealed a 47% frequency of naive CD4+ T-cells^28^, which have been shown to be enriched in hypermutated proviruses^57^. On average however, hypermutated proviruses comprised a median 19% (range 2%- 53%) of participants’ proviral pools, whereas those with packaging signal defects comprised a median 8% (range 4- 23%). Proviruses with inversions, gene scrambles, premature stop codons and HIV-human chimeras were uncommon, making up a median 2% (range 0.5%- 5%) of proviruses per participant.

As expected^50^, *nef* was the most commonly intact region in four participants (BC-001, BC-002, BC-004 and BC-027), and the second most commonly intact region in BC-003 and BC-021. The fraction of proviruses with an intact *nef* region ranged from 65% (127 sequences) for BC-027 to only 16% (115 sequences) for BC-003, due to the high proportion of hypermutated proviruses in the latter participant. The subset of *nef*-intact proviruses were those whose integration dates could be phylogenetically inferred, as described below.

### Proviral Clonality During Long-term ART

Consistent with the major role of clonal expansion in sustaining the HIV reservoir^21, 31– 33, 38–42, 44, 58^, a median 39% (range 17-55%) of proviruses were identical (100% sequence identity) to at least one other from that participant, where an average of 24 such “clonal sets” (range 16- 33) were recovered per participant (**Figure 2A**). BC-003 had the lowest proportion of clonal sequences, at 17% (n=121), while BC-027 had the highest, at 55% (n=107). BC-027 also harbored the most abundant clone: isolated 39 times, it harbored a ∼3,000 base deletion and made up 20% of the proviral pool (**Figure 2A**, bottom). BC-001’s three most frequent clones, which were recovered between 20-32 times each and included two sequences with packaging signal defects, together made up 24% of the proviral pool.

Clonal frequency significantly differed by proviral genomic integrity (**Figure 2B-E**). Across all participants, intact proviruses were nearly three times more likely to be part of a clonal set (19% of intact compared to 8% of other proviruses were clonal; Odds Ratio [OR] 2.7; p= 0.01, **Figure 2B**), as were proviruses with packaging signal defects (OR 2.8; p=0.0007; **Figure 2C**). Proviruses with large deletions were not preferentially clonal (p=0.6; **Figure 2D**), and hypermutated proviruses were less likely to be clonal (OR 0.51; p=0.0009, **Figure 2E**).

### Elucidating the ages of intact and defective proviruses and reservoir-origin viremia

We used a phylogenetic approach^13^ to estimate the ages of HIV sequences persisting on ART. To do this, we inferred within-host phylogenies relating longitudinal pre-ART plasma HIV RNA *nef* sequences with viral sequences sampled post-ART, whose integration dates we wished to estimate. The latter included near full-length proviruses for all participants, as well as QVOA outgrowth sequences, HIV RNA from on-ART viremia episodes and/or HIV RNA isolated after ART interruption for four participants. To mitigate the inherent uncertainty in within-host phylogenetic reconstruction, we inferred distributions of 1,500 - 4,500 phylogenies per participant using Bayesian approaches, and conditioned results over all trees.

We began by rooting each tree at the inferred most recent common ancestor (see methods). If plasma HIV RNA sampling goes back far enough, this root would represent the transmitted founder virus; if it does not go back far enough, it would represent a descendant of this founder. We then fit a linear model relating the root-to-tip genetic distances of unique pre-ART plasma HIV RNA *nef* sequences to their sampling dates. The slope of this line, which represents the average within-host pre-ART *nef* evolutionary rate, was then used to convert the root-to-tip distance of each post-ART sequence of interest to its integration date. We conditioned these dates over all trees that met our quality control criteria (see methods), yielding integration date point estimates and 95% highest posterior density (HPD) intervals for each sequence. As trees were inferred from *nef* sequences, only *nef*-intact proviruses could be dated: this amounted to a median 121 (range 70- 165) proviruses per participant, representing a median 32% (range 16-65%) of all proviruses collected.

#### Participant BC-001

Participant BC-001 was diagnosed with HIV in August 1996. ART was initiated in August 2006 but interrupted shortly thereafter, and durable viral suppression was not achieved until June 2008 (**Figure 3A**). We collected 102 plasma HIV RNA *nef* sequences from 15 pre-ART time points spanning 10 years, along with 317 proviruses sampled in June 2016, ∼9.5 years after ART initiation, 131 (41%) of which had an intact *nef* (**Figure 3A, Table 1**). We also isolated nine (five unique) HIV RNA *nef* sequences from plasma collected in September 2007, after ART was interrupted and viremia rebounded to 23,000 copies/mL, and one HIV RNA *nef* sequence from plasma collected in September 2015 when an isolated viremia “blip” to 76 copies/mL occurred. All 1,500 rooted within-host phylogenies showed strong molecular clock signal, yielding an average estimated *nef* pre-ART evolutionary rate of 3×10^−5^ (95% HPD 1.7×10^-5^ – 4.2×10^−5^) substitutions/nucleotide site/day, and a mean root date of February 1995 (95% HPD December 1993 – February 1996), approximately 18 months prior to the participant’s HIV diagnosis (example phylogeny and linear model in **Figures 3B and 3C**; amino acid highlighter plot in **Supplemental Figure 3A**). The phylogeny exhibited the “ladder-like” form typical of within-host HIV evolution, which is shaped by serial genetic bottlenecks imposed by host immune pressures^59–62^, a phenomenon that is apparent in the selective sweeps occurring at numerous Nef residues during untreated infection (**Supplemental Figure 3A**).

**Figure 3:**
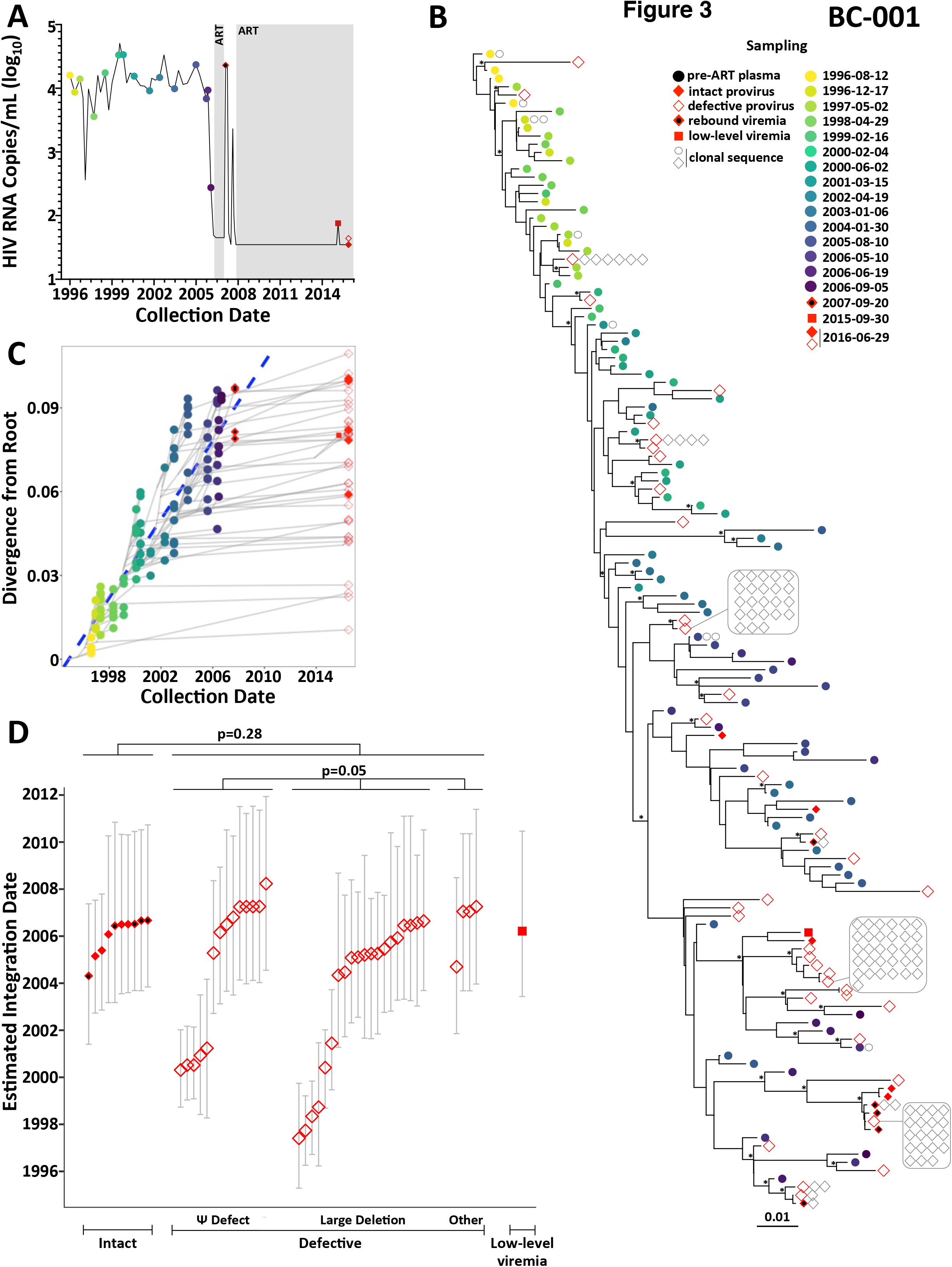
HIV evolu1onary reconstruc1on and on-ART sequence da1ng for par1cipant BC-001. *Panel A:* Clinical and sampling history. Plasma viral load is shown as a solid black line, with pre-ART plasma HIV RNA sampling dates shown as colored circles. Shaded periods denote ART. Red symbols denote sampling dates of on-ART sequences of interest, including intact proviruses (solid red diamonds), defective proviruses (open red diamonds), plasma rebound viremia following ART interruption (solid red diamond with black dot) and on-ART low-level viremia (solid red square). *Panel B:* Example within-host phylogeny, which is the highest likelihood tree derived from Bayesian inference, rooted at the most recent common ancestor as described in the methods. Scale in estimated substitutions per nucleotide site. Asterisks identify nodes supported by posterior probabilities ≥70%. *Panel C:* HIV sequence divergence- versus-time plot derived from the example phylogeny. The blue dashed line represents the linear model relating the root-to-Np distances of distinct pre-ART plasma HIV RNA sequences (colored circles) to their sampling times, which is used to convert the root- to-Np distances of distinct proviral sequences sampled during ART (red symbols) to their integration dates. Light grey lines trace the ancestral relationships between HIV sequences. Sequences from the last pre-ART timepoint were excluded from the linear model as the participant had initiated ART and viral load was decreasing at this time. *Panel D:* Integration date point estimates and 95% highest posterior density intervals for distinct post-ART sequences of interest, stratified by sequence type, that were derived from averaging results across all 1,500 passing trees for this participant. P-values compare the integration date point estimates between groups: the Mann-Whitney U test was used to compare the intact vs. combined defective categories (p=0.28), while the Kruskal-Wallis test was used to compare the different types of defective proviruses (p=0.05).

The unique HIV sequences sampled post-ART interspersed throughout the phylogeny, consistent with their continual archiving throughout infection. Averaging their phylogenetically-derived integration dates across all 1,500 trees revealed that the oldest of these sequences, a provirus with a large deletion, was estimated to have integrated in May 1997, 19 years prior to sampling, while the youngest was estimated to have integrated in March 2008, during the ART interruption, consistent with reservoir re-seeding during this rebound event. Despite this wide spread in integration dates, ∼50% of proviruses persisting during long-term ART dated to within 1.25 years of ART initiation (**Figure 3D**), and therefore harbored accumulated mutational adaptations to within-host pressures (**Supplemental Figure 3A**).

Notably, all of the old sequences sampled on ART (those estimated to have integrated in the first five years of infection, *i.e.* in ∼2001 or prior) were defective proviruses (**Figure 3D**). By contrast, all intact proviruses, as well as the HIV RNA sequences from the 2007 viremia rebound, were estimated to have integrated in the 2.75 years prior to ART. Though this is consistent with the hypothesis that intact HIV sequences persisting on ART are overall younger than the larger defective proviral pool, this comparison did not reach statistical significance (p= 0.28; **Figure 3D**), as many defective proviruses also dated to this period. Integration dates did not significantly differ between defective provirus types (Kruskal-Wallis p=0.05), though the very oldest all harbored large deletions. The single sequence isolated from the 2015 isolated viremia event dated to March 2006, close to ART initiation.

#### Participant BC-002

Participant BC-002 was diagnosed with HIV in April 1995, received non-suppressive dual ART between July 2000 and December 2006, after which viral suppression was finally achieved on triple ART (**Figure 4A**). Viral suppression was maintained until May 2011, after which frequent low-level viremia occurred, reaching a peak of 1,063 HIV RNA copies/mL in March 2013, despite no documented ART interruption during this time. We collected 160 plasma HIV RNA *nef* sequences from 17 time points spanning 9.5 years pre-ART (**Figure 4A, Table 1**), along with 265 proviruses sampled in August 2016, 9.6 years after ART initiation, 70 (26%) of which were *nef*-intact. We also isolated 13 (10 unique) plasma HIV RNA *nef* sequences from 2013 during on-ART viremia. All 1,500 within-host phylogenies exhibited strong molecular clock signal, yielding a mean pre-ART *nef* evolutionary rate of 1.5×10^−5^ (95% HPD 8.2×10^-6^ – 2.2×10^−5^) substitutions/site/day (example reconstruction in **Figures 4B, 4C**; highlighter plot in **Supplemental Figure 3B**) with a mean root date of May 1992 (95% HPD August 1989 – October 1994), three years prior to diagnosis. Proviral and HIV RNA sequences sampled on ART interspersed through the tree, where the oldest one, a defective provirus, was estimated to have integrated in the first year of infection, making it nearly 21 years old at time of sampling (**Figure 4D**). Nevertheless, ∼50% of sampled proviruses were estimated to have integrated in the 2.25 years prior to ART.

**Figure 4:**
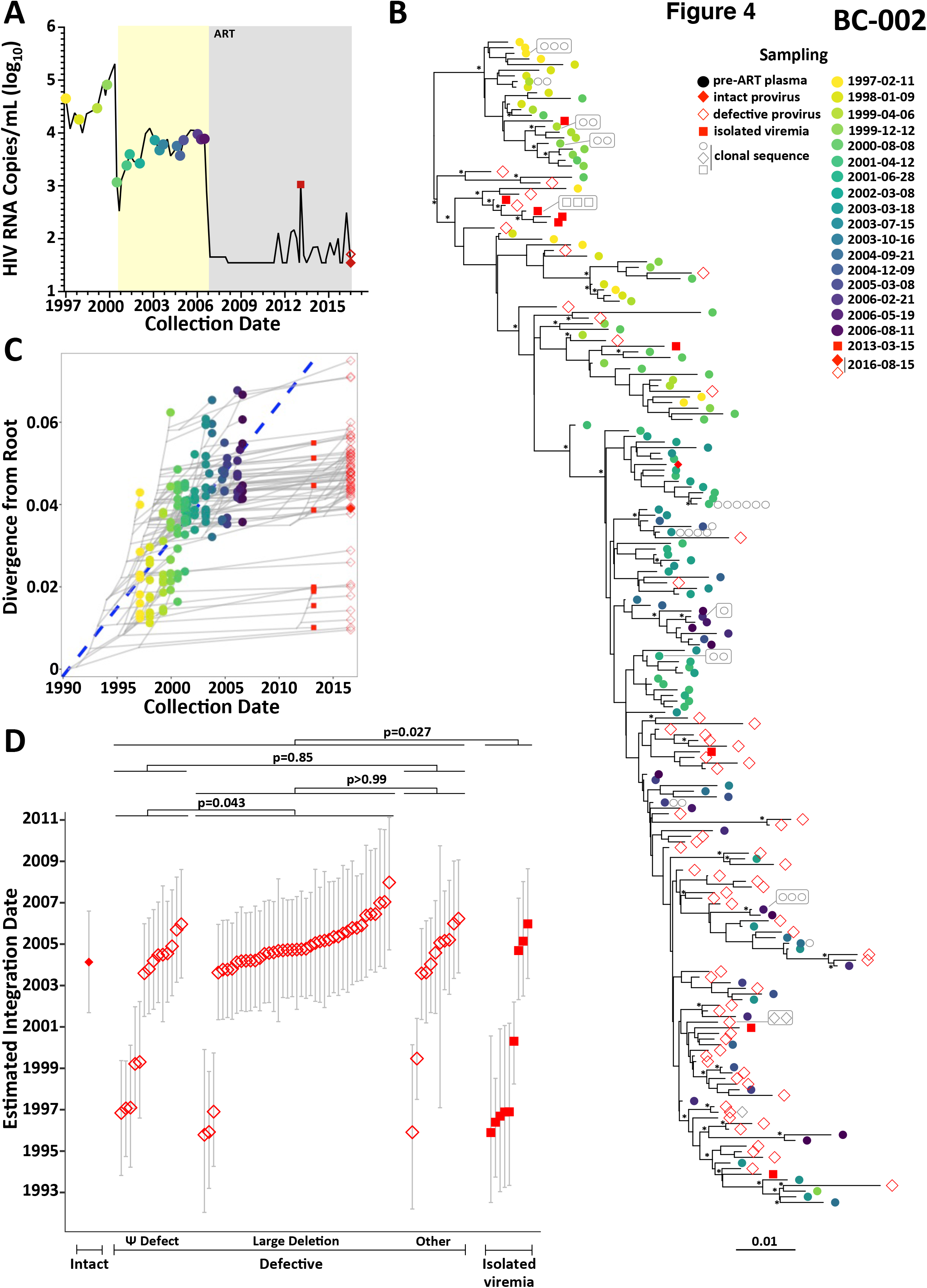
HIV evolu1onary reconstruc1on and on-ART sequence da1ng for par1cipant BC-002. Legend as in Figure 3, with the following modifications: In *panel A*, the yellow shading denotes the period of non-suppressive dual ART. In *panel D*, the Kruskal- Wallis test comparing the three types of defective proviruses was statistically significant, so the post-test p-values for the individual pairwise comparisons, corrected for mulNple comparisons, are shown.

Like BC-001, all of BC-002’s old proviruses (those dating to before the initiation of dual ART in July 2000) were defective. By contrast, the single intact provirus was estimated to have integrated in 2004, though many defective proviruses also dated to around this time. Notably, the HIV RNA sequences recovered during the persistent on-ART viremia period were genetically diverse and included sequences dating as far back as 1995; in fact, the sequences that emerged in plasma during this 2013 event were on average older than sampled proviruses (p=0.027; **Figure 4D**). Four of these sequences, one of which was recovered four times, mapped to a single subclade near the top of the tree, where the most closely related sequence was a provirus whose sole defect was a premature stop codon in Vif (**Figure 4B**). This further supports the recent finding that defective proviruses can contribute to low-level viremia on ART^52^, though we cannot rule out an unsampled intact provirus as the origin. Proviruses with large deletions were slightly older than those with Ψ defects (p= 0.043) though otherwise no significant age differences were found between defective proviral types (**Figure 4D**).

#### Participant BC-003

Participant BC-003 was diagnosed with HIV in 2002. HIV was suppressed on ART in October 2007 and largely maintained for ∼ 9 years, except for isolated low-level viremia (<250 HIV RNA copies/mL) in April 2010 and June 2015, from which HIV amplification was unsuccessful (**Figure 5A**). We isolated 122 plasma HIV RNA *nef* sequences from 8 pre-ART time points spanning 4.75 years, along with 733 proviruses on ART, of which 115 (16%) were *nef*-intact (**Figure 5A, Table 1**). We also isolated one full and one partial HIV RNA genome from limiting-dilution QVOA performed on the same sample as the proviral isolation. All 4,500 within-host phylogenies showed strong molecular clock signal, yielding a mean pre-ART *nef* estimated evolutionary rate of 6.8×10^−5^ (95% HPD 4×10^−5^ – 1×10^−4^) substitutions/site/day and a mean root date of December 2001 (95% HPD February 2001 – August 2002), which was only a few months prior to diagnosis (example reconstruction in **Figure 5B, 5C;** highlighter plot in **Supplemental Figure 3C**). Overall, BC-003’s proviral pool was markedly skewed in age: all but two proviruses dated to the 2.75 years prior to ART (**Figure 5C, 5D**). Nevertheless, intact proviruses and QVOA outgrowth viruses were on average significantly younger than defective proviruses (p=0.03; **Figure 5D**). Though defective proviral types did not overall differ in age (Kruskal-Wallis p= 0.8), the oldest proviruses all harbored large deletions.

**Figure 5:**
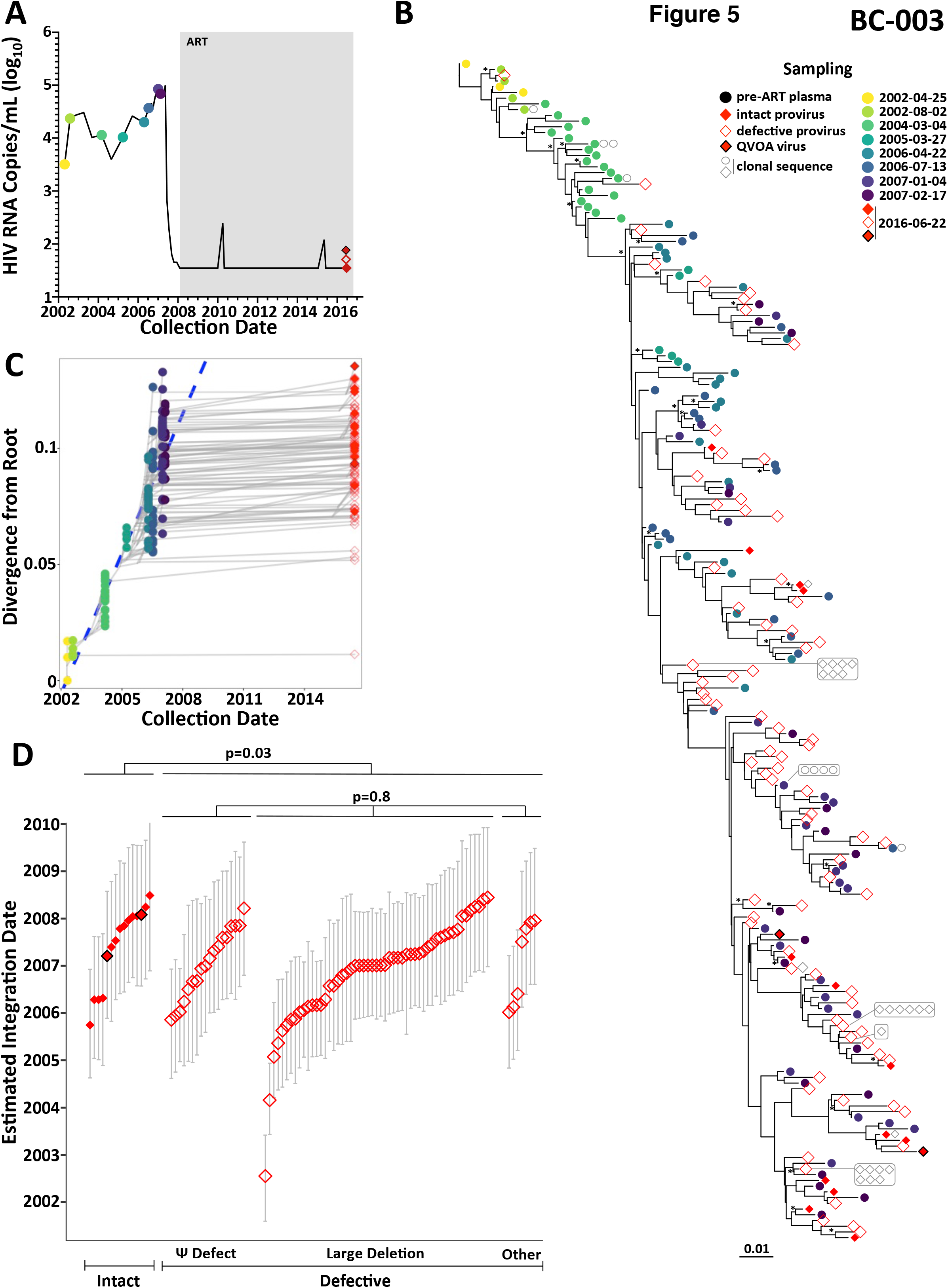
HIV evolu1onary reconstruc1on and on-ART sequence da1ng for par1cipant BC-003. Legend as in Figure 3, with the following modificaNons: QVOA outgrowth virus sequences are denoted with solid red diamonds with a black outline.

#### Participant BC-004

Participant BC-004 initiated ART less than two years following diagnosis, which was much earlier than the other participants (**Figure 6A**). Intermittent low-level viremia occurred in the first four years of ART, but suppression was maintained thereafter except for an isolated measurement of 3,120 copies/mL in 2019. There was no documented ART interruption at this time, and antiretroviral resistance genotyping predicted that all drugs retained full activity. We isolated 65 plasma HIV RNA *nef* sequences from 4 pre-ART time points spanning 8 months, along with 440 proviral genomes (165; 38% *nef*-intact), and 7 unique HIV RNA genomes from QVOA in July 2016 (**Figure 6A, Table 1**). We also isolated 46 (4 unique) and 44 (4 unique) plasma HIV RNA *nef* sequences from the on-ART viremia in 2011 and 2019. Of the 1,500 rooted within-host phylogenies, only 379 (25%) had sufficient molecular clock signal to pass quality control, which is not surprising given that early ART initiation limits within-host HIV evolution^63–67^. The passing trees yielded a mean pre-ART *nef* evolutionary rate of 1×10^−4^ (95% HPD 4.9×10^−5^ – 1.6×10^−4^) substitutions/site/day (example reconstruction in **Figures 6B, 6C;** highlighter plot in **Supplemental Figure 3D**) and a mean root date of April 2005 (95% HPD October 2004 – September 2005). This, combined with the limited viral diversity in this individual (**Supplemental Figure 3D**), is consistent with HIV diagnosis during early infection. Due to the inherent uncertainty in phylogenies inferred from limited-diversity datasets, estimated integration dates for this participant have wide 95% HPD intervals, and should be cautiously interpreted. Overall, BC-004’s intact and defective proviruses did not differ in terms of age (p=0.41), nor did defective proviruses differ in age based on defect type (p=0.48) (**Figure 6D**).

**Figure 6:**
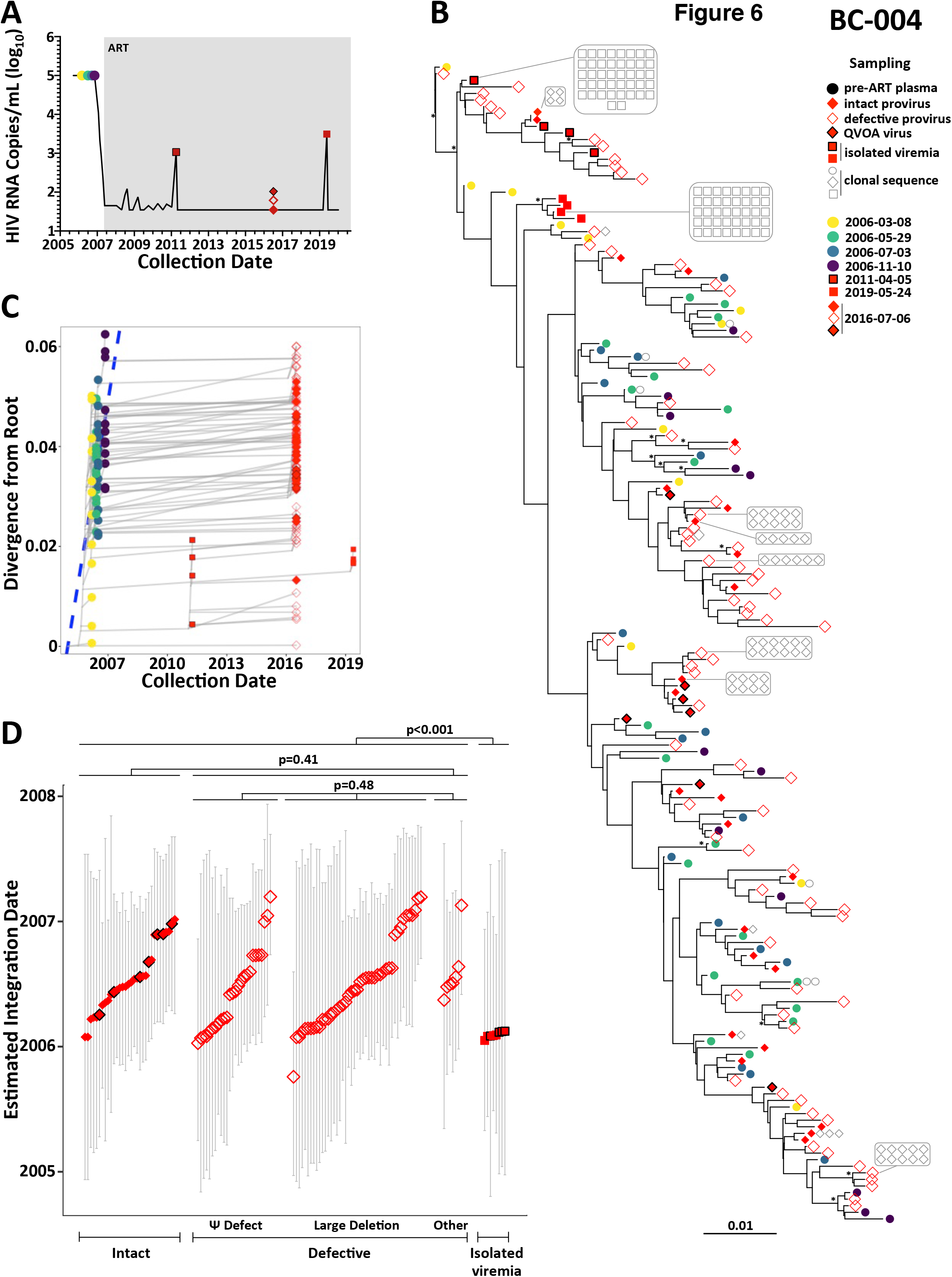
HIV evolu1onary reconstruc1on and on-ART sequence da1ng for par1cipant BC-004. Legend as in Figure 3, with the following modifications: QVOA outgrowth virus sequences are denoted with solid red diamonds with a black outline. Unique symbols are used to differentiate the 2011 (solid red square with black outline) and 2019 (solid red square) on-ART viremia events.

Nevertheless, the oldest provirus, estimated to have integrated in October 2005, six months following the estimated transmission date, was defective due to a large deletion (**Figure 6D**). Moreover, sequences isolated from the on-ART viremia in 2011 and 2019 exclusively dated to early infection (January/February 2006), and in fact were on average older than sampled proviruses (p<0.001) (**Figure 6D**). The 2011 viremia sequences, one of which was recovered 43 times, all fell within a single diverse subclade that featured both intact and defective proviruses sampled on ART (**Figure 6B**, near top of tree), while the 2019 viremia sequences, one of which was recovered 41 times, were more restricted in terms of diversity and formed an exclusive subcluster also relatively near the root (**Figure 6B**).

#### Participant BC-021

Participant BC-021 was diagnosed with HIV in November 2002 and achieved viral suppression on ART in April 2007 (**Figure 7A**). Suppression was largely maintained except for low-level viremia to 367 HIV RNA copies/mL in May 2018 from which sequence isolation was unsuccessful. We isolated 221 plasma HIV RNA *nef* sequences from 8 pre-ART time points spanning 2.5 years, and 386 proviruses on ART in July 2019, of which 93 (24%) were *nef*-intact (**Figure 7A, Table 1**). All 3,000 rooted phylogenies demonstrated strong molecular clock signal, yielding a mean pre-ART *nef* evolutionary rate of 8.4×10^−5^ (95% HPD 5.1×10^−5^ -1.2 ×10^−4^) substitutions/site/day (example reconstruction in **Figure 7B, 7C**; highlighter plot in **Supplemental Figure 3E**) and a mean root date of April 2002 (95% HPD December 2001- August 2002), seven months prior to diagnosis. BC-021’s on-ART proviral pool was markedly skewed in age, with all but two unique sequences dating to the 2 years prior to ART (**Figure 7D**). Though intact and defective proviruses did not differ in age (p=0.99), the only two old proviruses, which dated to 2003, 16 years prior to sampling, both harbored large deletions (p=0.1 for comparison with other defective proviral types, **Figure 7D**).

**Figure 7:**
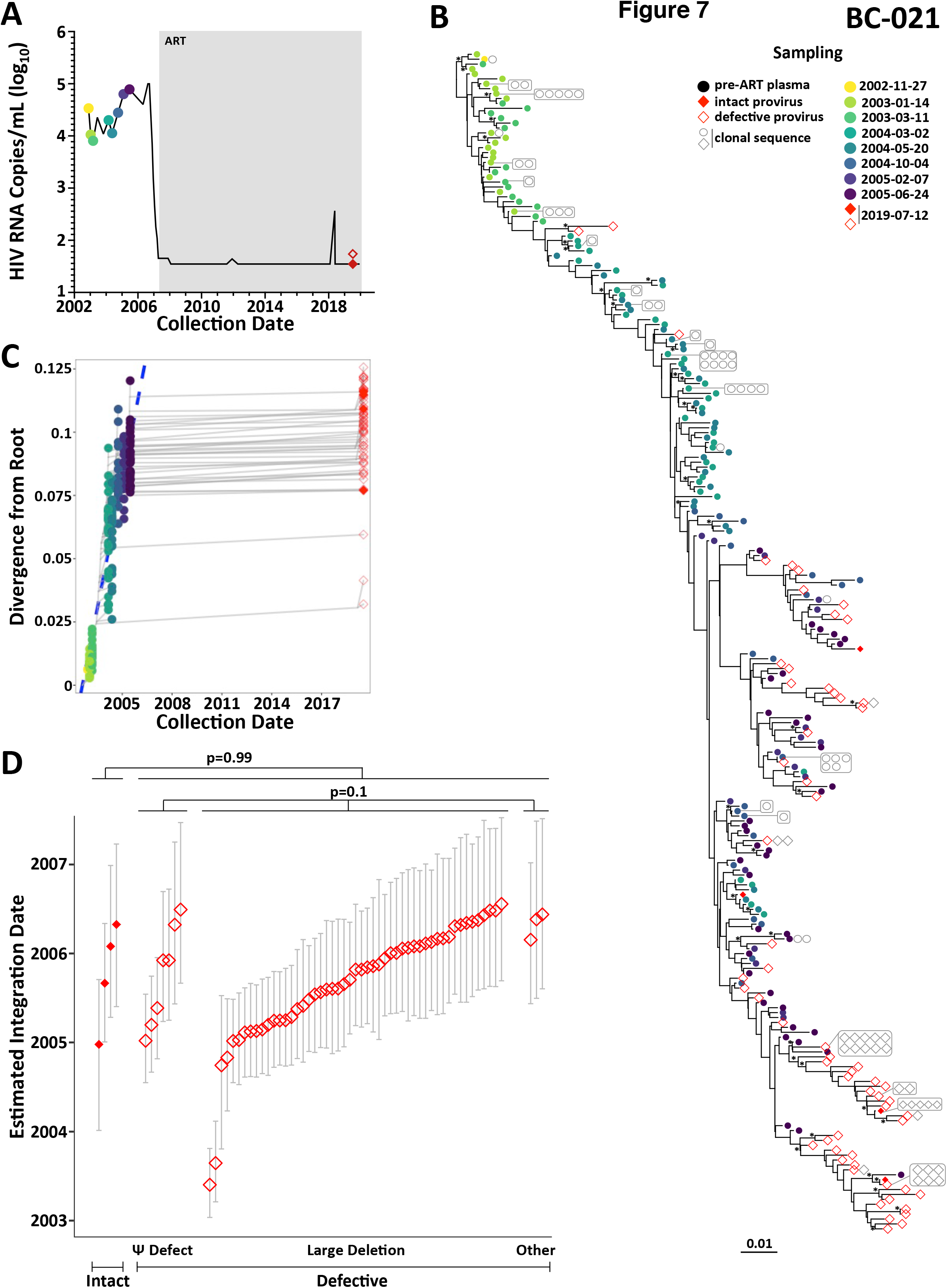
HIV evolu1onary reconstruc1on and on-ART sequence da1ng for par1cipant BC-021. Legend as in Figure 3.

#### Participant BC-027

Participant BC-027 was diagnosed with HIV in 1985, received various non-suppressive ART regimens beginning in the mid-1990s, and finally achieved viral suppression in December 2011 (**Figure 8A**). Suppression was maintained except for two low-level viremia events <80 HIV RNA copies/mL from which sequence isolation was unsuccessful. We isolated 215 pre-ART plasma HIV RNA *nef* sequences from 7 time points spanning a 14.25 year period, along with 195 proviruses on-ART in 2019, of which 127 (65%) were *nef*-intact (**Figure 8A, Table 1**). All 1,500 within-host rooted phylogenies displayed strong molecular clock signal, yielding a mean (95% HPD) pre-ART *nef* estimated evolutionary rate of 1.5×10^−5^ (9.1×10^-6^ – 2.1×10^−5^) substitutions/site/day (example reconstruction in **Figure 8B, 8C**; highlighter plot in **Supplemental Figure 3F**). The mean root date was September 1992 (95% HPD January 1990 – March 1995), indicating that we did not reconstruct back to the founder virus but rather to one of its descendants. Though integration dates did not differ between intact and defective proviruses (p=0.96, **Figure 8D**), all of the old proviruses, the oldest of which dated to 1996, 23 years prior to sampling, were defective. BC-027 was therefore the only participant for whom no proviruses dating to the earliest years of infection were recovered (1996 was > 10 years after diagnosis).

**Figure 8:**
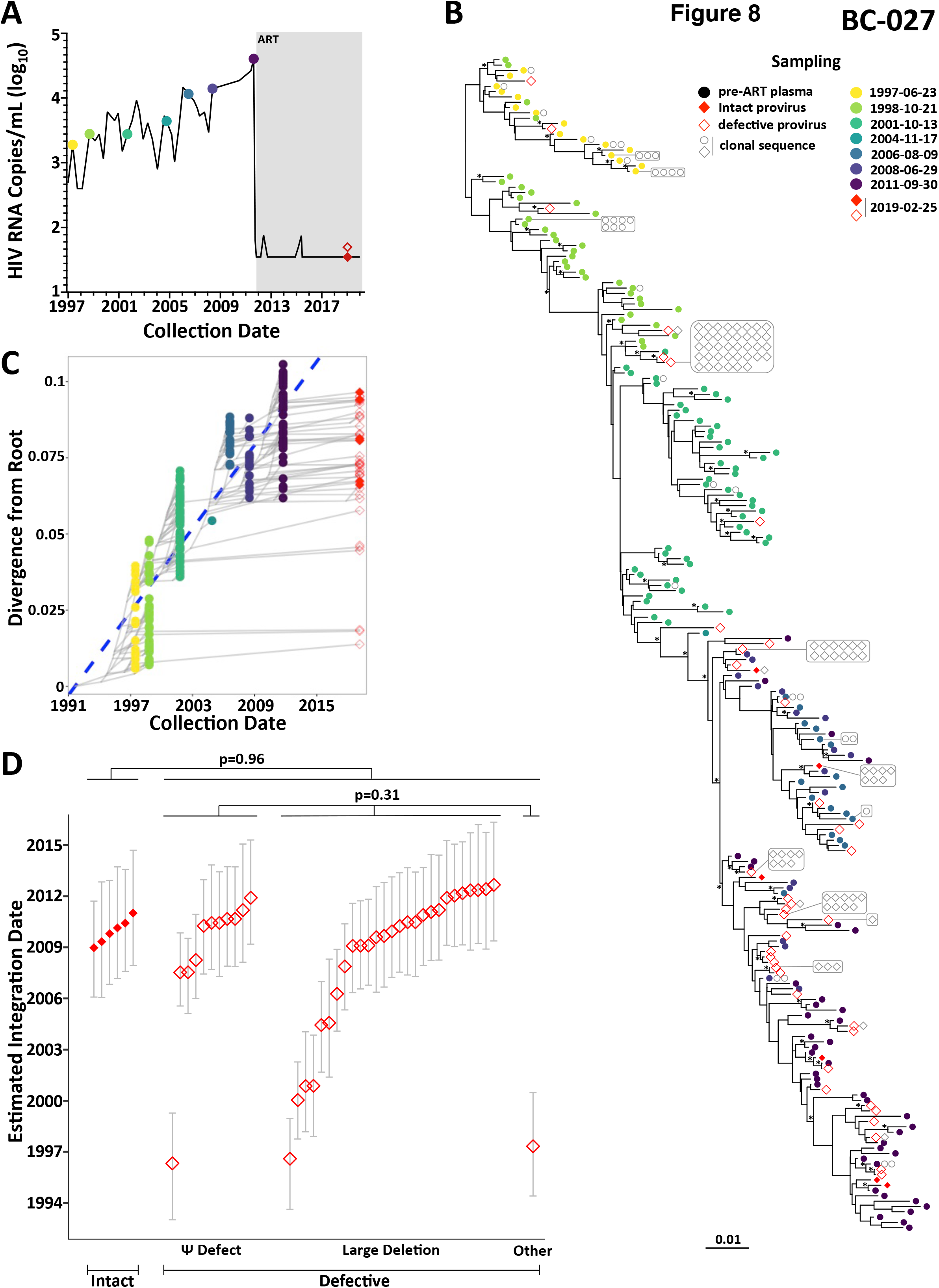
HIV evolu1onary reconstruc1on and on-ART sequence da1ng for par1cipant BC-027. Legend as in Figure 3.

Defective proviruses did not differ significantly in age (Kruskal-Wallis p=0.31) (**Figure 8D**).

### Clones do not differ from unique sequences in terms of age

We investigated whether clonal sequences differed from unique ones in terms of their overall age distribution. Here, members of a clonal set were collapsed into a single data point for analysis, and sequences from plasma viral rebound and on-ART viremia were not considered, as only *nef* was sequenced and therefore clonality cannot be established. We observed no significant differences in age distribution between clonal and unique sequences in any participant (**Supplementary Figure 4**, all p>0.1). This remained true when sequences were stratified by genomic integrity (all p>0.12).

## Discussion

Reservoir dynamics studies should distinguish intact proviruses from the vast background of defective ones, as only the former can re-seed infection if ART is stopped. We therefore resampled near-full-length proviruses persisting on ART, along with reservoir-origin HIV RNA, from four participants for whom only subgenomic proviral sequencing had previously been performed (BC-001 through BC-004)^13, 28^, along with two others. This allowed us to explore the relationship between proviral genomic integrity and longevity, as well as the within-host origins of plasma viremia. As expected, given that proviruses were sampled after an average of more than 9 years on ART^48^, only 4% were intact, with some participants (*e.g.* BC-002) having as few as 0.4% intact. Proviruses with large deletions typically dominated, where, consistent with previous studies^38, 48^, we identified shared HIV genomic breakpoints within and across individuals that illuminate how such deletions reproducibly occur.

Our observation that intact and Ψ-defective proviruses were nearly three times more likely to be clonal, while hypermutated proviruses were twofold less likely, extend our understanding of clonal expansion in reservoir maintenance^21, 31, 38–42, 44, 45^. While these differences in part reflect cellular distribution^57^ (intact and Ψ-defective proviruses tend to be enriched in effector memory CD4+ T-cells^39, 44, 57, 68^, which have the highest proliferative capacity^69^, while hypermutated proviruses tend to be enriched in naive CD4+ T-cells^39, 44, 57, 68, 70^, which have the lowest proliferative capacity^69^), they are also consistent with intact and Ψ-defective proviruses being more dependent on clonal expansion for survival. This is because they are at higher risk of elimination by cytopathic effects or immune responses after reactivation (at least some Ψ- defective proviruses can produce HIV proteins^71^ and even virions^72^). As not all members of a clone will reactivate upon stimulation^73, 74^, expansion enhances the likelihood that at least some members will persist^75^. In contrast, grossly defective proviruses rely less on clonal expansion for survival, as their risk of elimination is inherently lower due to limited or no viral antigen presentation^21, 53^.

Also consistent with prior studies^14–16, 29^, participants’ on-ART proviral pools ranged from modestly (*e.g*. BC-001) to substantially (*e.g* BC-003 and BC-021) skewed towards viral variants archived in the years immediately preceding ART. This is consistent with continual reservoir seeding - and turnover - during untreated infection, such that, if ART is not initiated until chronic infection, many ancestral within-host lineages will have already been eliminated by this time^14, 17, 18^. Nevertheless, and consistent with prior studies that sampled subgenomic proviral sequences on-ART^13–16, 28^, we recovered at least one sequence dating to early infection in all participants but BC-027 (this individual’s oldest provirus, though 23 years old when sampled, dated to more than 10 years after diagnosis). Importantly, the oldest recovered proviruses were exclusively defective, usually due to large deletions, whereas all intact proviruses dated to within approximately 3 years preceding ART. This indicates that intact proviruses have shorter lifespans than those with gross defects. This is consistent with their more rapid decay during ART^21–23, 25, 26^, and likely during untreated infection as well^14, 17, 18^, though the latter has not been explicitly demonstrated.

The observation that the oldest on-ART proviruses are exclusively defective also resolves a discordance in the literature. It explains why studies that recovered subgenomic proviral sequences on ART (which are largely defective) routinely recovered proviruses dating to early infection^13–16, 28^, whereas the study that exclusively dated *ex vivo* viral outgrowth sequences sampled on ART yielded hardly any sequences dating to this time^29^. The latter study nevertheless did find a handful of “old” intact viruses, whereas we found none (except in BC-004, for whom all proviruses dated to the short period between infection and ART). This difference may be because both the time to ART initiation (except for BC-004) and the time on ART were substantially longer in the present study, allowing more time for intact proviruses to be eliminated *in vivo*. Taken together with existing data^40, 41, 44^, our findings indicate that intact proviruses are at a survival disadvantage compared to their grossly defective counterparts and are thereby more dependent on clonal expansion for persistence.

Our observations also suggest that low-level/isolated viremia on ART can have distinct within-host origins from rebound viremia. Though the sequence recovered from BC-001’s low-level viremia dated to the year before ART, those recovered from BC-002 and BC-004’s on-ART viremia indicated that proviruses capable of *reactivating* to produce viremia can be genetically heterogeneous, and can originate from very ancestral proviruses, which may in some cases be defective. The existence of genetically defective virions is supported by their presence during untreated infection^76^ (though recently-infected, rather than reservoir cells, would be the likely source during that time), and the recent discovery that low-level viremia can originate from defective proviruses^51, 72^. By contrast, HIV RNA rebounding in plasma after ART interruption dated to the years just prior to ART (BC-001), consistent with its origin from intact proviruses dating to that same period.

Our observation that intact proviruses (and rebound viremia) exclusively dated to the later years of untreated infection has implications for immune-based cure strategies, because sequences from this period will have substantially adapted to within-host selective pressures (**Supplemental Figure 3** illustrates the selective sweeps that typify this process^67, 77^). Indeed, Human Leukocyte Antigen (HLA) class I-restricted escape mutations were apparent in the five participants who initiated ART in advanced chronic infection; these included the C*06:02-restricted Nef-125H adaptation^78^ at position 6 of the C*06-restricted-YT9 epitope (Nef 120- 128)^79, 80^ in BC-001, the C*07:01-restricted Nef-105Q adaptation^78^ at position 1 of the C*07- restricted KY11 epitope (Nef 120-128)^78^ in BC-003, the B*35:01-restricted Nef-81F adaptation^78^ at the C-terminus of the B*35:01-restricted VY8 epitope (Nef 74-81)^81^ in BC-027, and others.

Studies of rebound HIV from larger numbers of individuals in a within-host evolutionary context will help establish whether rebound sequences are on average even more adapted to host immune responses than the overall intact proviral pool. If so, this would be consistent with the notion that rebound is a selective process, where the viruses that first appear in plasma at high levels are not necessarily those that reactivated first, but rather those that host immune responses, particularly antibodies, subsequently fail to control^82, 83^ (this may also explain why *in vivo* rebound and *in vitro* outgrowth HIV don’t always match^84^).

Our study has some limitations. Despite extensive sampling, we recovered fewer intact proviruses than predicted based on the IPDA, reflecting the inefficiency of long-range PCR^56^. As we did not determine integration site, we cannot definitively state that identical sequences are clonal, though the likelihood is high as all proviruses were sequenced end-to-end, and a prior study demonstrated that proviruses with 100% sequence identity also share integration sites^85^.

We also acknowledge that sequences isolated only once may still be part of a clonal set^86^. HIV integration dates are derived from a model that assumes a strict molecular clock, which may not be ideal over long time frames^87^. Nevertheless, the observation that the oldest proviruses are defective is apparent even without the model, as these were always the closest to the root of the phylogeny. As we could only “date” *nef*-intact proviruses, we cannot rule out that *nef*-defective proviruses have a different age distribution, which is possible as *nef* may promote proviral longevity^57^. Nevertheless, studies that have used *gag* and *env* for dating^14–16, 29^ have produced similar proviral age distributions, suggesting that our results are not overly biased. There are also no ideal solutions to this, as the abundance of large deletions means that no single HIV region can be used to date all proviruses phylogenetically, nor can hypermutated sequences be dated using such approaches.

Despite these limitations, our study is the first to compare the ages of defective and intact proviruses on ART, along with reservoir-origin HIV RNA, in context of within-host HIV evolutionary history. As participants were sampled during the slower second phase of on-ART decay^22, 24^, recovered proviruses represent truly long-lived populations. We addressed within-host phylogenetic reconstruction uncertainty by inferring 1,500- 4,500 trees per participant. Rather than using clustering approaches, which can only “date” sequences of interest to the specific time points when plasma HIV RNA was sampled pre-ART^29, 88^, our approach allowed us to date sequences to before (or after) pre-ART plasma sampling. This was particularly important for BC- 021, whose pre-ART sampling stops ∼2 years prior to ART, but where substantial HIV evolution and proviral archiving undoubtedly took place during this time.

In summary, the oldest proviruses persisting during long-term ART were exclusively genetically defective, whereas intact proviruses and rebound HIV RNA dated nearer to ART initiation and were thus enriched in mutations consistent with accumulated adaptation to host pressures. Intact proviruses were also more likely to be clonal. This indicates that genomic integrity shortens a provirus’ lifespan, likely due to increased risk of viral reactivation, antigen production and elimination^49, 53, 71^, such that intact proviruses are significantly more dependent on clonal expansion for survival than their defective counterparts. By contrast, on-ART viremia sometimes belonged to very old and possibly defective within-host lineages^72^, underscoring the need to better understand the biological and clinical management implications of the large burden of defective proviruses that persist during ART. Nevertheless, our results provide further evidence that cure strategies will need to eliminate an intact viral reservoir that is clonally- enriched, genetically “younger”, and thus more adapted to its host.

## Methods

### Participants and Sampling

We recruited six participants living with HIV. All were male, and had initiated triple ART a median 11 (range 2.2- 26.5) years after estimated HIV infection. At the time of peripheral blood mononuclear cell (PBMC) sampling on ART, plasma viral loads had been largely suppressed for a median 9.3 (range 7.2- 12.2) years. A median 8 (range 4- 17) pre-ART longitudinal plasma samples per participant were used to reconstruct within-host HIV evolutionary histories. We also studied plasma from a viremia rebound event following ART interruption in one participant (BC-001), and plasma from on-ART low-level and/or isolated viremia events in three participants (BC-001; BC-002; BC-004). Human Leukocyte Antigen (HLA) Class I typing was performed on DNA extracted from whole blood or CD4+ T-cells, using a sequence-based method^89^.

### Ethics statement

Ethical approval to conduct this study was obtained from the research ethics boards of Providence Health Care/ University of British Columbia and Simon Fraser University. All participants provided written informed consent.

### Amplification and Sequencing of Plasma HIV RNA

HIV RNA *nef* sequences were isolated from longitudinal pre-ART plasma samples, rebound viremia and on-ART viremia events as follows. Total nucleic acids were extracted from 500mL of plasma on the NucliSENS EasyMag (bioMerieux), and subjected to DNAse treatment if the original plasma viral load was < 250 HIV RNA copies/mL. cDNA, generated using HIV-specific primers, was endpoint-diluted such that subsequent nested PCR reactions yielded no more than 30% positive amplicons, as previously described^13, 17^. Amplicons were sequenced using a 3130xl or 3730xl Automated DNA Sequencer (Applied BioSystems) and chromatograms were analyzed using Sequencher (v.5.0, Gene Codes). Sequences with nucleotide mixtures, hypermutation or suspected within-host recombination (identified using RDP4^90^) were removed prior to phylogenetic inference. HIV drug resistance genotyping was performed on select on-ART viremia samples using standard approaches^91^, and interpretations were performed using the Stanford HIV drug resistance database^92^.

### Intact Proviral DNA Assay (IPDA)

Intact and total proviral DNA was quantified in CD4+ T-cells isolated by negative selection from on-ART PBMC using the Intact Proviral DNA Assay (IPDA)^50, 93^, as previously described^93^, where XhoI restriction enzyme (New England Biolabs) was added to each ddPCR reaction to aid in droplet formation per the manufacturer’s recommendation. For participant BC- 004, an autologous *env* probe (VIC-CCTTGGGTT**<u>TC</u>**TGGGA-MGBNFQ) was used, as the published IPDA probe failed due to these polymorphisms^93^.

### Near full-length HIV Proviral Amplification and Sequencing

Single-template, near full-length HIV proviral amplification was performed on genomic DNA extracted from CD4+ T-cells using Platinum Taq DNA Polymerase High Fidelity (Invitrogen), where IPDA-determined total proviral loads were used to dilute DNA such that no more than 30% of resulting nested PCR reactions yielded an amplicon (protocol described in ^40^). Amplicons were sequenced (Illumina MiSeq) and reads were *de novo* assembled using an in-house modification of the Iterative Virus Assembler^94^ (IVA) implemented in the custom software MiCall (http://github.com/cfe-lab/MiCall) to generate a consensus sequence. Each sequence was verified to have been sequenced end-to-end (by locating at least part of the 2nd round PCR primers at both 5’ and 3’ ends) and to have a minimum read depth of ≥100 over all positions.

Sequences not meeting these criteria were discarded. We validated our pipeline by single-genome sequencing the provirus integrated within the J-Lat 9.2 cell line^95^ in 146 independent replicates, as well as by sequencing a panel of nine in-house engineered pNL4-3 plasmids in which we had deleted large HIV genomic regions^96–99^. The J-Lat validation yielded 25 consensus base errors out of a total 1,375,174 bases sequenced (*i.e.* an error rate of 1.8 x 10^−5^), while the plasmid validation correctly reconstructed the deletion breakpoints 100% of the time. The genomic integrity of sequenced proviruses was determined using an in-house modification of the open-source software HIV SeqinR^100^, where an intact classification required all HIV reading frames, including accessory proteins, to be intact. Intactness classifications for all non-hypermutated proviruses longer than 8,000bp were manually confirmed by checking each HIV gene for the presence of start and stop codons (where applicable), internal stop codons, hypermutation, or other defects, and checking for known defects in the packaging signal region^38^. Sequences with 100% identity across the entire amplicon were considered identical and clonal.

### Sequence isolation from Quantitative Viral Outgrowth Assay (QVOA)

QVOA^101^ was performed for four participants for whom sufficient biological material was available (BC-001, BC-002, BC-003 and BC-004), as previously described^93^. Only two of these participants, BC-003 and BC-004, yielded viral outgrowth. For these, near-full length HIV RNA genomes were RT-PCR amplified from viral culture supernatants in five overlapping amplicons, sequenced on an Illumina MiSeq, and assembled as described above.

### Within-host HIV evolutionary reconstruction and phylogenetic dating

Within-host phylogenies were inferred from *nef* sequence alignments comprising all unique plasma, proviral and QVOA sequences collected per participant. To mitigate uncertainty in within-host HIV phylogenetic reconstruction, we inferred a median 1,500 (range 1,500- 4,500) phylogenies per participant using Bayesian approaches. To do this, the best-fitting substitution models for each dataset was first determined using jModelTest (version 2.1.10 ^102^). Markov Chain Monte Carlo (MCMC) methods were then used to infer a random sample of phylogenies for each participant. Two parallel runs with MCMC chains of five million generations each, sampled every 10,000 generations, were performed in Mr. Bayes (version 3.2.5 ^103^), employing the best-fitting nucleotide substitution model (or second best fitting model when runs did not readily converge) and model-specific or default priors. Convergence was assessed by ensuring the deviation of split frequencies was <0.03, the effective sample size of all parameters was ≥200 and through visual inspection of parameter traces in Tracer (version1.7.2, ^104^). The first 25% of generations were discarded as burn-in. Nodal support values are derived from Bayesian posterior probabilities from the consensus tree.

The integration dates of on-ART sequences of interest were subsequently inferred using a phylogenetic approach^13^ as follows. First, each participant’s phylogenies were rooted at the estimated most recent common ancestor, by identifying the location that maximized the correlation between the root-to-tip distance and the sampling date of the pre-ART plasma HIV RNA sequences (as within-host sequence divergence from the transmitted/founder virus increases over time^87, 105, 106^). Linear regression was then used to relate the sampling dates of the pre-ART plasma HIV RNA sequences and their root-to-tip divergence, where the slope of the regression represents the average within-host *nef* evolutionary rate and the x-intercept represents the root date. We assessed model fit by comparing the Akaike Information Criterion (AIC) of the model to that of the null model (a zero slope), where a ΔAIC ≥10 was required to pass quality control. Passing linear models were then used to convert root-to-tip distances of on-ART sequences of interest to their estimated integration dates, which were then averaged across all passing models per participant. Highest posterior density intervals were computed using the R package HDInterval (version 0.2.2).

Between-host phylogenies shown in Supplemental Figures 1 and 2 were inferred using *gag* and *nef* sequences under a maximum likelihood model. Briefly, sequences were aligned in a codon-aware manner in MAFFT (version 7.475 ^107^) and manually edited in AliView (version 1.19 ^108^). Phylogenies were inferred with IQ-Tree^109^ following automated model selection using ModelFinder^110^ with an AIC selection criterion. Branch support values were derived from 1,000 bootstraps. Phylogenies were visualized using the R package ggtree^111^.

### Statistical Analyses

Unless otherwise indicated, statistics were computed using GraphPad Prism 8 or R version 4.2.1 implemented in RStudio v2021-01-06 or greater.

## Data Availability

Previously published sequences used in this study have the following accession numbers: MG822917, MG822918, MG822920-MG822961, MG822963- MG823015, MN600002- MN600010, MG823016, MG823017, MG823019-MG823143, MN600011-MN600053, MN600054- MN600175 and MN600183- MN600247. GenBank accession numbers for the HIV RNA *nef* sequences from pre-ART plasma and rebound and on-ART viremia collected in this study are: OQ723958- OQ7244 and OQ750829- OQ750838. GenBank accession numbers for defective HIV proviral sequences are: OQ750839- OQ753083, while those for intact HIV proviruses and QVOA outgrowth viruses are pending.

## Funding Statement and Acknowledgements

We are grateful to the study participants, without whom this research would not be possible. We thank Gursev Anmole and Harrison Omondi for helpful discussion.

This work was supported in part by the Canadian Institutes for Health Research (CIHR) through a project grant (PJT-159625 to ZLB and JBJ) and a team grant (HB1-164063 to ZLB and MAB). This work was also supported in part by the National Institutes of Health (NIH) through the Martin Delaney “REACH’ Collaboratory (NIH grant 1-UM1AI164565-01 to ZLB, MAB and RBJ), which is supported by the following NIH cofounding institutes: NIMH, NIDA, NINDS, NIDDK, NHLBI and NIAID. NNK is supported by a CIHR Vanier Canada Graduate Scholarship. AS and BRJ are supported by CIHR CGS Doctoral Awards. HS is supported by a CIHR MSc award. EB was supported by an Undergraduate Student Research Award from Simon Fraser University. ZLB was supported by a Scholar Award from Michael Smith Health Research BC. The content is solely the responsibility of the authors and does not necessarily represent the official views of the National Institutes of Health or other funders.

## Author Contributions

NNK and ZLB conceived and designed the study. NNK, WD, DK, CJB, DM, TMM, HS and EB performed experiments and collected data. NNK, AS, GQL and BRJ analyzed data. MH and CJB provided sample access. RBJ, MAB and JBJ provided critical feedback and expertise to study design. NNK and ZLB wrote the manuscript. All authors critically reviewed the manuscript.

