## Supplemental Figures (1-4) and Table (S1) for "HIV reservoirs are dominated by genetically younger and clonally enriched proviruses"

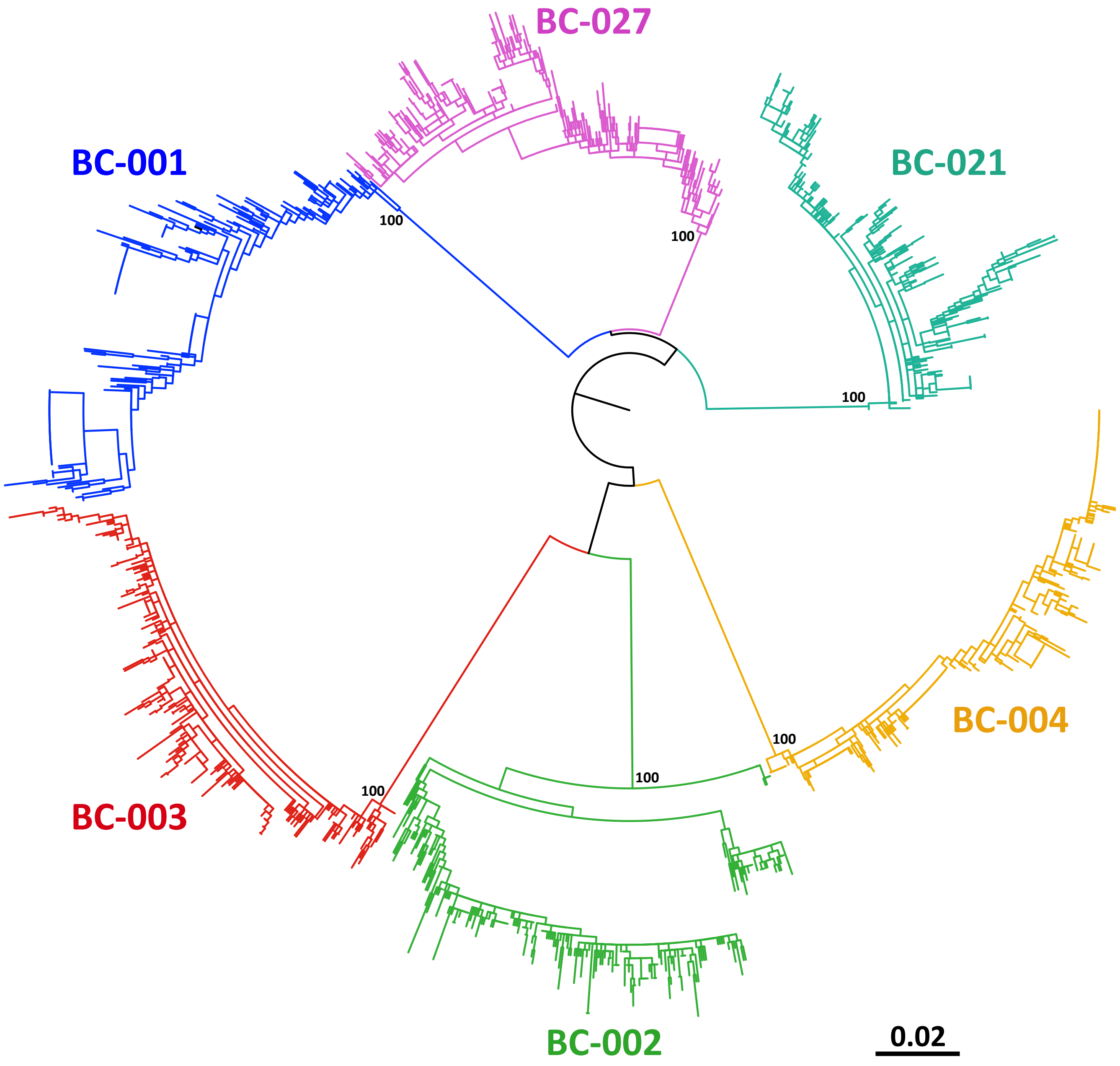


**Figure S1: Between-host phylogeny inferred from *nef* sequence alignments.** Maximum-likelihood phylogeny inferred from all pre-ART plasma HIV RNA sequences along with all on-ART sequences of interest with an intact *nef* gene. Phylogeny is mid-point rooted. Numbers on internal branches indicate bootstrap values. Scale in estimated substitutions per nucleotide site.

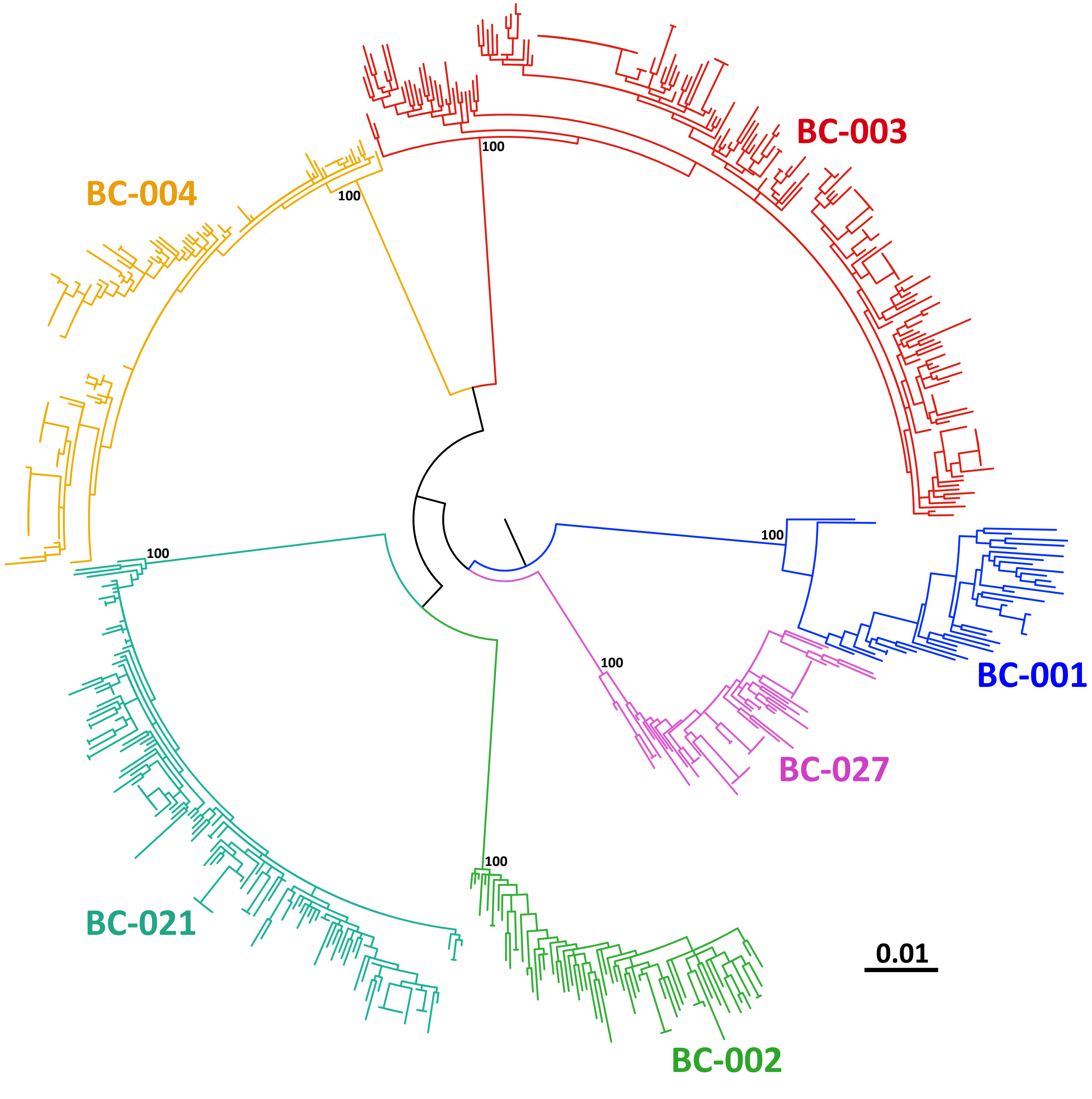


**Figure S2: Between-host phylogeny inferred from *gag* sequence alignments.** Same as Figure S1, but inferred from all on-ART proviral sequences with an intact *gag* gene.

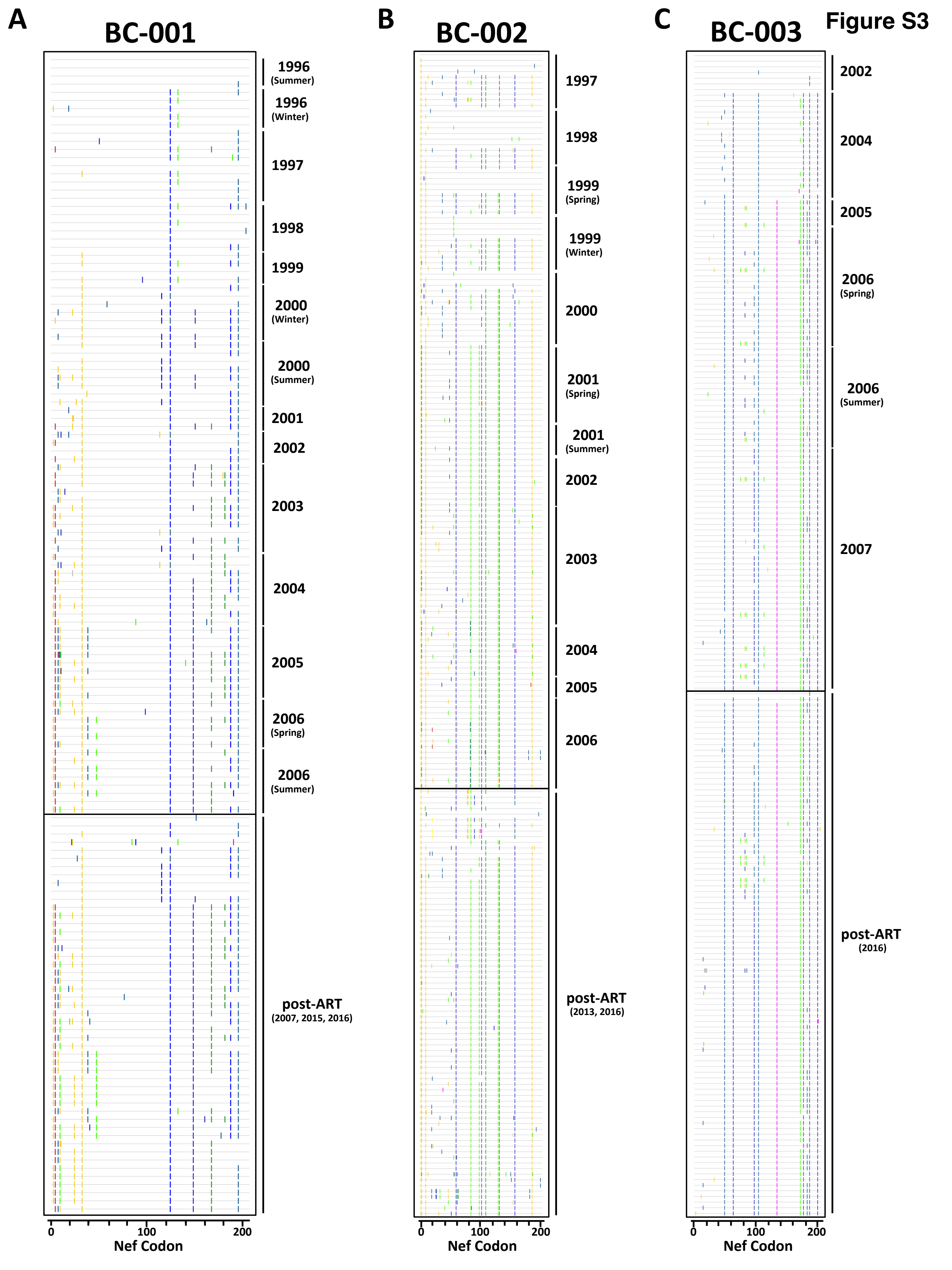
Figure S3
A
BC-001
1996 (Summer)
1996 (Winter)
1997
1998
1999
2000 (Winter)
2000 (Summer)
2001
2002
2003
2004
2005
2006 (Spring)
2006 (Summer)
post-ART (2007, 2015, 2016)
0
100
200
Nef Codon
B
BC-002
1997
1998
1999 (Spring)
1999 (Winter)
2000
2001 (Spring)
2001 (Summer)
2002
2003
2004
2005
2006
post-ART (2013, 2016)
0
100
200
Nef Codon
C
BC-003
2002
2004
2005
2006 (Spring)
2006 (Summer)
2007
post-ART (2016)
0
100
200
Nef Codon

Figure S3

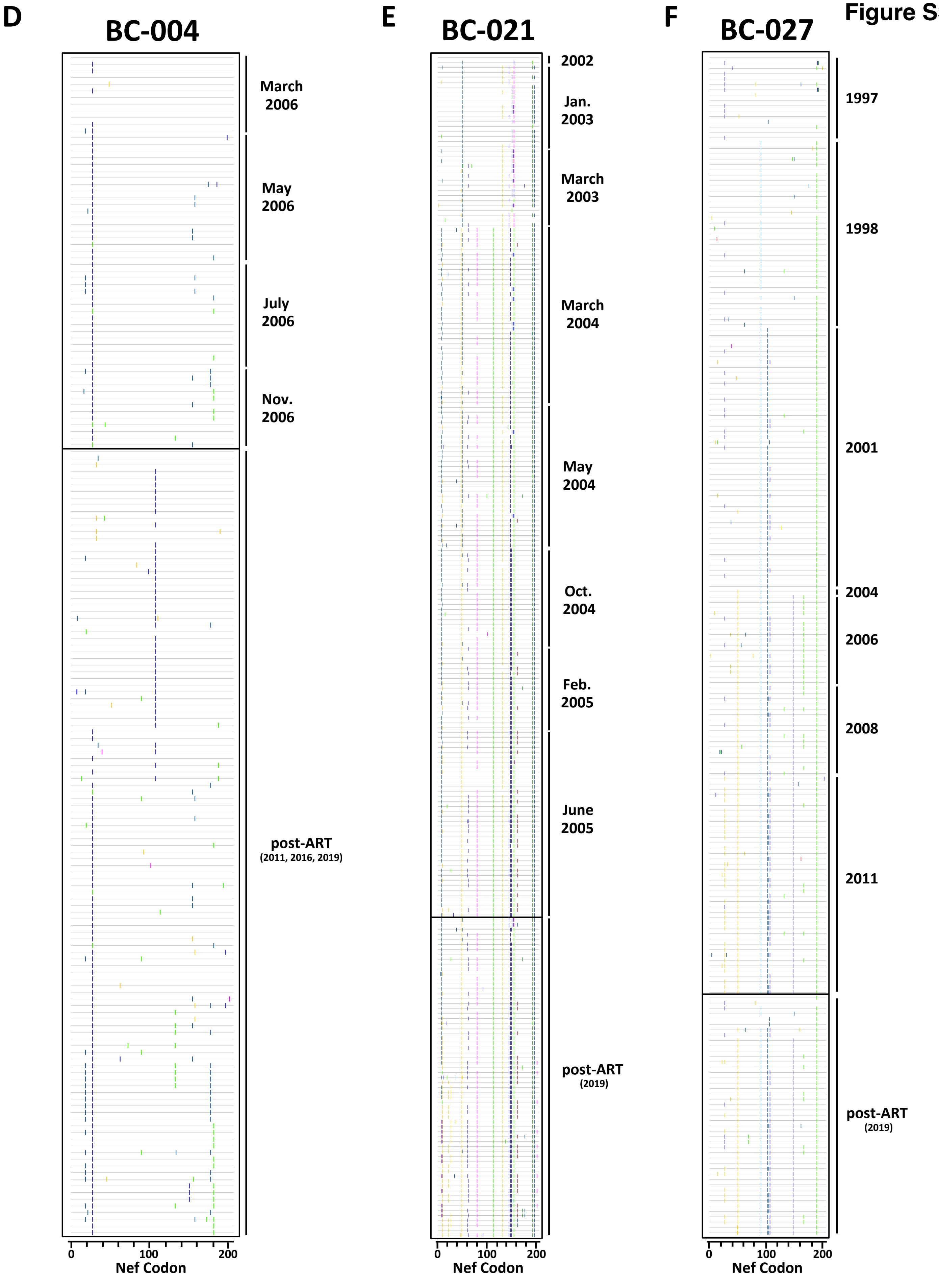
D
BC-004
March 2006
May 2006
July 2006
Nov. 2006
post-ART
(2011, 2016, 2019)
0
100
200
Nef Codon
E
BC-021
2002
Jan. 2003
March 2003
March 2004
May 2004
Oct. 2004
Feb. 2005
June 2005
post-ART
(2019)
0
100
200
Nef Codon
F
BC-027
1997
1998
2001
2004
2006
2008
2011
post-ART
(2019)
0
100
200
Nef Codon

**Figure S3: Amino acid highlighter plots depicting pre-ART *nef* evolution and post-ART *nef* sequence composition.** *panel A: nef* amino acid highlighter plot of BC-001's pre-ART plasma and post-ART sequences. The reference (top) sequence represents the pre-ART plasma sequence closest to the root of the participant's example phylogeny (shown in Figure 3), where colored ticks in sequences beneath this denote non-synonymous substitutions relative to this reference. Pre-ART *nef* sequences are ordered according to their sampling date, while post-ART *nef* sequences are ordered according to their inferred integration date. Note the frequent selection of amino acid changes that then "sweep" through all subsequent pre-ART plasma sequences, and that are preserved in sequences persisting post-ART. *panels B-F:* Highlighter plots for participants BC-002, BC-003, BC-004, BC-021 and BC-027, respectively.

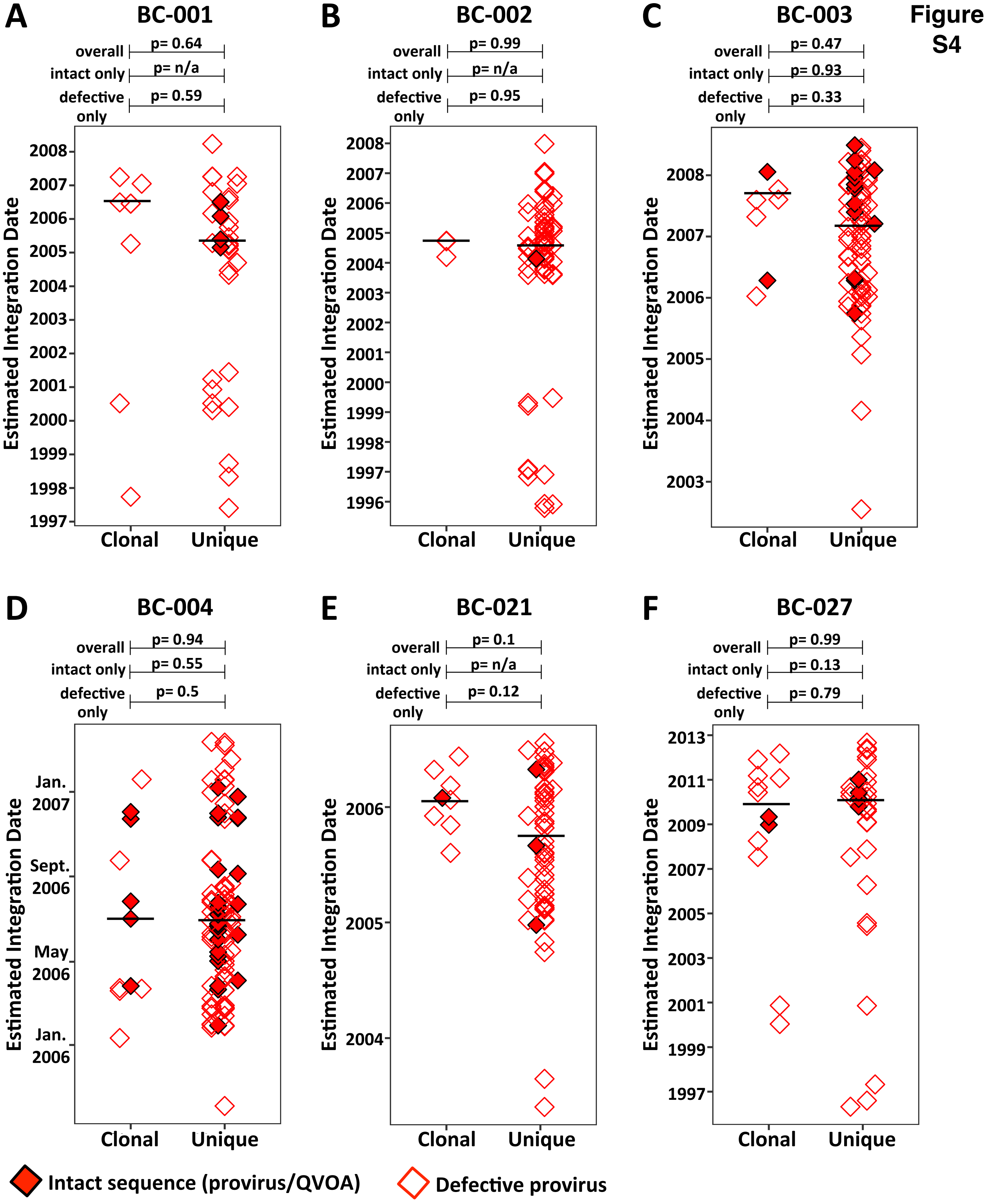
Figure S4
A
BC-001
overall p= 0.64
intact only p= n/a
defective only p= 0.59
B
BC-002
overall p= 0.99
intact only p= n/a
defective only p= 0.95
C
BC-003
overall p= 0.47
intact only p= 0.93
defective only p= 0.33
D
BC-004
overall p= 0.94
intact only p= 0.55
defective only p= 0.5
Jan. 2007
Sept. 2006
May 2006
Jan. 2006
E
BC-021
overall p= 0.1
intact only p= n/a
defective only p= 0.12
F
BC-027
overall p= 0.99
intact only p= 0.13
defective only p= 0.79
Estimated Integration Date
Clonal
Unique
Intact sequence (provirus/QVOA)
Defective provirus

**Figure S4: Lack of relationship between sequence clonality and age.** Estimated integration dates of distinct proviruses that were observed more than once (clonal) versus only once (unique), stratified by participant. Open diamonds denote defective proviruses; red diamonds with black outline include intact proviruses and HIV RNA sequences recovered from *ex vivo* reactivation (QVOA), if any. Black lines indicate median estimated integration date for each group. For the comparisons indicated above the plots, p-values were determined using the Mann-Whitney test and were not corrected for multiple comparisons.

**Supplemental Table 1: Proviral Integrity and Reservoir Size**

| | BC-001 | BC-002 | BC-003 | BC-004 | BC-021 | BC-027 | Total |
|---|---|---|---|---|---|---|---|
| Total proviral sequences (n) | 317 | 265 | 733 | 440 | 386 | 195 | 2,336 |
| Intact | 5 | 1 | 15 | 47 | 9 | 14 | 91 |
| Total defective | 312 | 264 | 718 | 393 | 377 | 181 | 2,245 |
| Hypermutated | 95 | 34 | 386 | 107 | 7 | 7 | 636 |
| Ψ defective | 68 | 15 | 34 | 46 | 15 | 44 | 222 |
| Large deletion | 138 | 201 | 285 | 227 | 349 | 129 | 1,329 |
| Inversion | 4 | 9 | 3 | 7 | 2 | 1 | 26 |
| Premature stop | 2 | 4 | 6 | 4 | 3 | - | 19 |
| Scramble | 1 | 1 | 3 | - | 1 | - | 6 |
| HIV-Human chimera | 4 | - | 1 | 2 | - | - | 7 |
| Total HIV copies per million CD4+ T-cells | 1,757 | 4,052 | 1,930 | 1,851 | 765 | 518 | |
| Intact copies | 85 | 199 | 238 | 160 | 70 | 32 | |
| Defective copies | 1,672 | 3,853 | 1,692 | 1,691 | 695 | 486 | |
